# Diagnostic Potential of Shallow Depth Gut Metagenomics Sequencing for Atherosclerotic Cardiovascular Disease Risk Stratification

**DOI:** 10.1101/2023.10.02.560614

**Authors:** Justin C. Sing, Owen Whitley, Sachin Davis

## Abstract

**Background:** Cardiovascular disease, specifically atherosclerotic cardiovascular disease (ACVD), presents a burden on society in terms of financial resources and quality of life that is only expected to grow in the coming years. Current diagnostic methods lack the sensitivity or specificity to confidently screen for atherosclerotic cardiovascular disease and are either costly, invasive, or both. Building on previous research linking ACVD to changes in the microbiome, we hypothesized that one could build a machine learning classifier for ACVD that makes use of shallow depth microbiome data.

**Methods and Findings:** Raw metagenomics Illumina paired end sequencing data was downloaded from the European Bioinformatics Institute (EBI) public database under accession ERP023788. The dataset includes 383 Han Chinese subjects of which 170 are control subjects and 214 are ACVD subjects. The raw sequencing data was subsampled using seqtk to generate *in silico* shallow depth datasets at various depths. Each subsampled experiment was processed through KneadData for quality control, and finally relative abundances were acquired through Kraken2. Based on *in-silico* down sampling experiments, we demonstrate that species level information is still captured at read depths as low as 50K reads per sample, consistent with current literature. In addition, the shallow depth data (50K) contains relevant taxa information for stratification of ACVD patients. Differential expression analysis identified a variety of *Streptococcus* species and *Escherichia coli* enriched in ACVD patients and a few low abundance *Bacteroides* species in ACVD patients, which have been previously reported.

**Conclusion:** Here, using publicly available data, we show with *in silico* experiments that species-level microbiome information is preserved at low sequencing depths that would allow a screening or diagnostic tool to be cost-competitive with currently used methods such as stress ECGs and stress ECHO tests. Additionally, we show that we can indeed make a microbiome-based model with performance metrics comparable to or better than front-line or mid-tier tests.

## Introduction

Cardiovascular disease (CVD) is one of the leading causes of death worldwide with an estimated mortality rate of 17.9 million people each year, of which 655,000 of these people are from the United States alone. ^1,2^ It is estimated that approximated 75% of CVD deaths occur in low to middle income countries, and there has been increasing evidence of rising CVD incidences in previously declining high-income countries. ^3^ CVD is not only an ongoing public health issue resulting in a high mortality rate, but also incurs rising health care and lost productivity costs. ^3^ The United States is predicted to have 40% of its population diagnosed with CVD by 2030, resulting in an increase in total costs from $445 billion USD in 2010 to $1094 billion USD by 2030, a 71% increase. ^4,5^ Similarly, CVD could cost Europeans €170 billion each year, and Canadians $22 billion CAD each year. ^5-7^ However, the inflating healthcare costs have shown value as life expectancy has improved following healthcare intervention, albeit there is still room for improvement in bringing costs down while still effectively ameliorating cardiovascular health.

The gut microbiome plays an integral role in human health but conversely also contributes to pathogenesis of disease. ^8^ Although the gut microbiome is defined as the community of bacteria, viruses, protozoa and fungi present in the gastrointestinal tract (GIT), the microbe-host interplay extends beyond the gut influencing the metabolic, immune, and neuroendocrine systems. ^9^ There have been several studies investigating the connection between CVD and the gut microbiome, with increasing evidence of correlations between microbiome composition and CVD suggesting dysbiosis of the gut microbiome contributes to or is caused by pathogenesis of CVD.^10^

Over the past decade different Next Generation Sequencing methodologies, such as PCR amplicon sequencing, metagenomics sequencing, metatranscriptomic sequencing and viromic sequencing have been developed to investigate the gut microbiome. ^11^ Amplicon sequencing has been widely adopted for investigating bacterial and fungal community composition, however, this method only allows the targeting of a single gene and also introduces bias towards the specific primer sequence used. ^11^ On the other hand, metagenomics ‘shotgun’ sequencing stochastically samples the metagenome, enabling a more detailed view of the structural and functional aspects of the gut microbiome. ^11^ Metatranscriptomics allows for a greater depth of insight into the gut microbiome, in that it reports microbe and cell activity through a snapshot of the microbial gene expression profile. ^11^ Although metatranscriptomics might give deeper insight into functionally active microbes, it is costly especially for large-scale studies. In contrast, while metagenomics can also have a high price point, it is possible to reduce the cost by sequencing at a shallower depth. ^12^

The data revolution has garnered much interest in software and algorithmic development in artificial intelligence (AI) within the health care sector aiding clinical decision support and managing electronic health records. ^13^ Furthermore, it is perceived that AI will also have a positive impact on precision medicine, as it becomes possible to track a patient’s health state over time (i.e., recurrent medical visits) allowing concurrent disease counselling as well as early onset anomaly detection of disease. ^13^ Specifically in the context of CVD, AI has shown to be fruitful in disease risk stratification, disease management and therapeutic intervention using clinical and laboratory tests, echocardiography and imaging data. ^13,14^ However, some of these tests are expensive (e.g. electrocardiogram stress tests and computed tomography angiograms, costing between $200 and $500 USD), and require the patients to make several visits to the clinician or hospital to get a prognosis.Here, we show the feasibility of using shallow depth metagenomics sequencing, as low as 50K reads, based on *in-silico* subsampling of publicly available data (Jie et. al. ^15^, 55.2 million reads per sample) to predict for CVD.

Here, we show the feasibility of using shallow depth metagenomics sequencing, as low as 50K reads, based on *in-silico* subsampling of publicly available data (Jie and colleagues, 2017 ^15^, 55.2 million reads per sample) to predict for CVD.

## Results

### Evaluation of Effect of Sequencing Depth on Feature Detection

In order to evaluate the potential for shallow sequencing as a means of obtaining high information content for lower cost, we ran a series of *in silico* experiments to assess preservation of information at lower sequencing depths. To ensure the randomness of *in silico* subsampling, we performed subsampling at a depth of 10K reads per sample, cleaning data and performing abundance estimation in a pipeline illustrated in Figure 1 (see *Online Methods* for more details). Based on the in-silico seed subsampling experiments we verified that across the different seed subsampling experiments, the distribution of total non-zero (i.e. non-zero abundance in a sample) and overlapping (between different runs in identical samples) features identified was similar across the six 10K reads subsampled datasets (Figure S1). Briefly, seed numbers are used to allow different subsamples of a given dataset in a reproducible manner, with each seed number generating an identical subsample from run to run. Subsequently, to determine the minimum reasonable depth at which information is preserved with respect to deep metagenomic sequencing, we ran the same computational pipeline at a variety of simulated sequencing depths.

**Figure 1:**
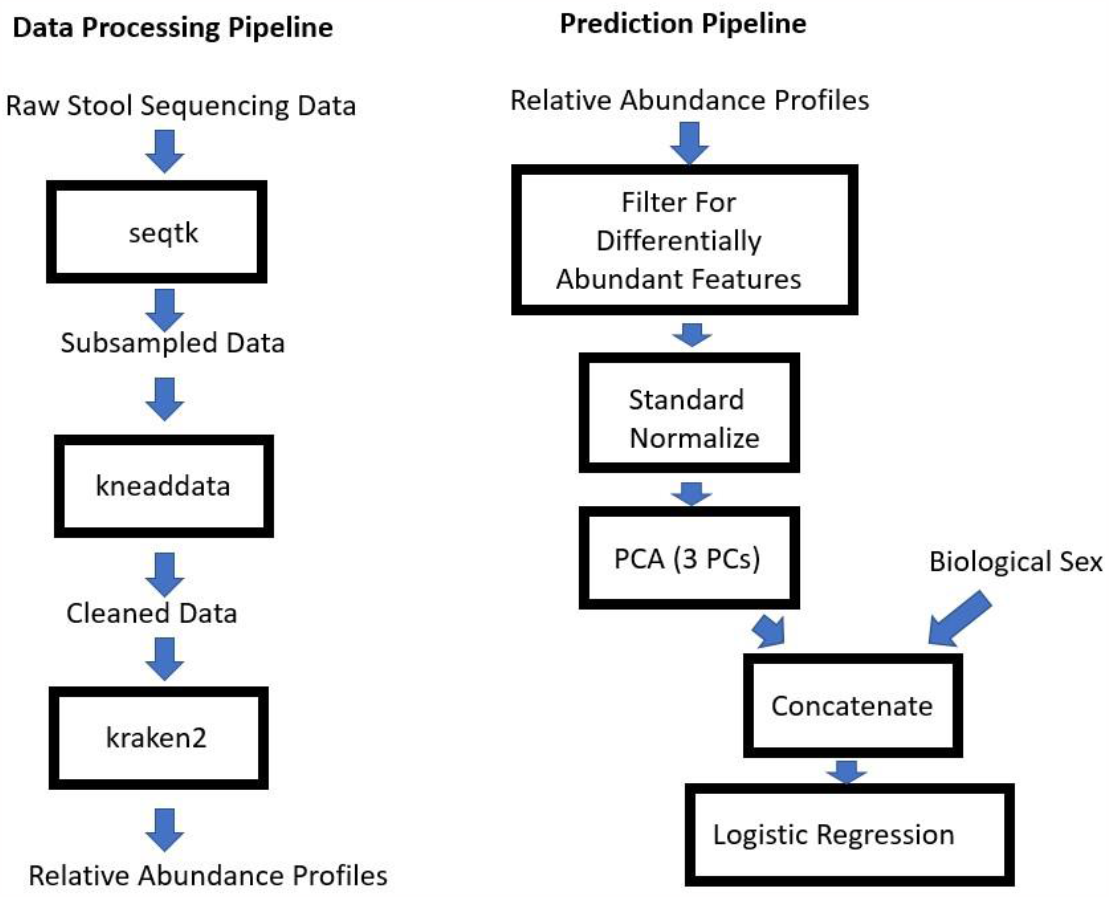
Overview of Data Processing and Prediction Workflow. Left: Data processing pipeline to generate subsampled, cleaned data. Right: Prediction pipeline. See *Online Methods* for full details.

We compared features detected at read depths ranging from 500 to 5 million to those detected at 10 million with a variety of metrics, including number of features detected, overlap of features, and correlation of abundance of overlapping features (Figure 2). Consistent with previously published research^12^, correlation of features detected at lower depths with those detected at 10 million reads was high down to 5K (Figure 2D). While this is somewhat lower than reported in Hillman and colleagues’ analysis ^12^, our results are consistent with the notion that relative abundance information can be preserved well below 1 million reads. We noted that with 10K and lower reads, the number of features overlapping with those detected at 10 million reads dropped off sharply. Thus, we elected to select 50K depth for further analyses involving microbiome species abundance profiles. Overall, our results demonstrate the preservation of species level microbiome information (i.e. features detected and relative abundance) at low sequencing depths, making shallow sequencing a suitable data generation process for a machine learning classifier.

**Figure 2:**
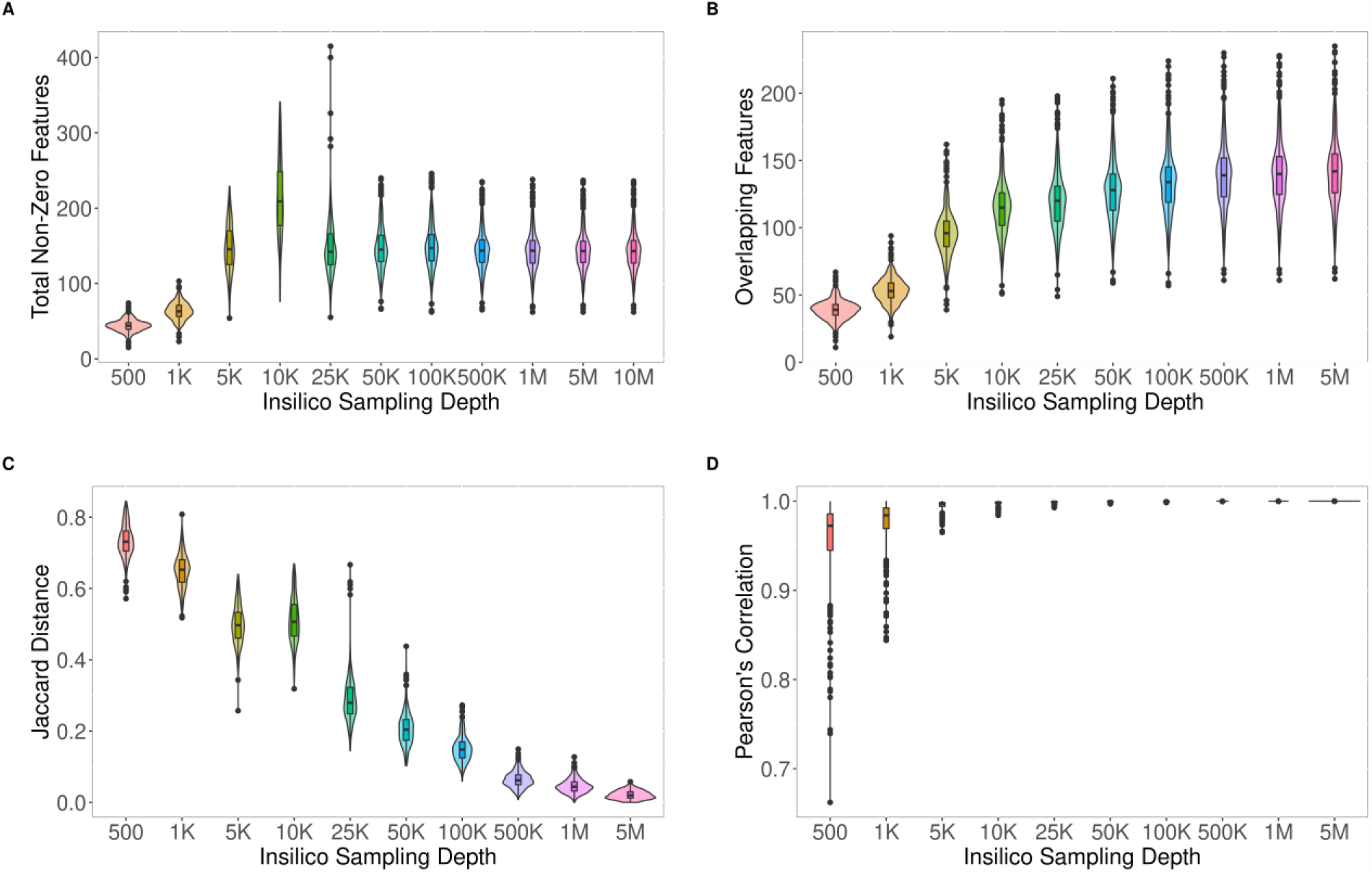
Subsampling Preserves Species Abundance Information at 10K Depth. For subsampling experiments, all 384 samples were subjected to subsampling as described in Figure 1 at the specified depth. The ‘reference’ depth that all samples were compared to was 10 million reads. **(A)** Total detected features (non-zero proportion) in data. **(B)** Number of overlapping features between sequences at a given depth and features detected at 10M depth. **(C)** Jaccard distance between features detected at a given depth and those detected at 10M depth **(D)** Pearson correlation of overlapping features between samples subsampled at varying depth and their 10M depth counterparts.

### Exploration of Features for Model Development

In order to assess if microbial features could potentially distinguish between ACVD patients and healthy patients, we ran a differential abundance analysis. We noticed that at different taxonomic levels (i.e. class and species) there were indeed differentially abundant microbial taxa (1% FDR) (Figure 3A, B, Table 3). To translate differential abundance analysis into a feature selection tool, we wrote a portion of our classification pipeline to intake a microbial abundance profile, perform differential abundance analysis, filter features for those passing an FDR threshold of 5% for the ACVD positive and negative classes, followed by principal component analysis (PCA) dimensionality reduction on that feature set. While not all ACVD positive patients could be distinguished from ACVD negative patients, the principal components appeared to separate a large portion of these populations (Figure 3C, D).

**Table 2.** Fisher’s Exact Test and Contingency Table for Gender and ACVD Association.

| <u>Contingency Table</u> |  |  |
| --- | --- | --- |
|  | Female | Male |
| Health Cohort | 101 | 68 |
| ACVD Cohort | 53 | 157 |
| <u>Fisher's Exact Test</u> |  |  |
| Odd's Ratio | 4.34 |  |
| P-Value | 7.75E-12 |  |

**Table 3.**
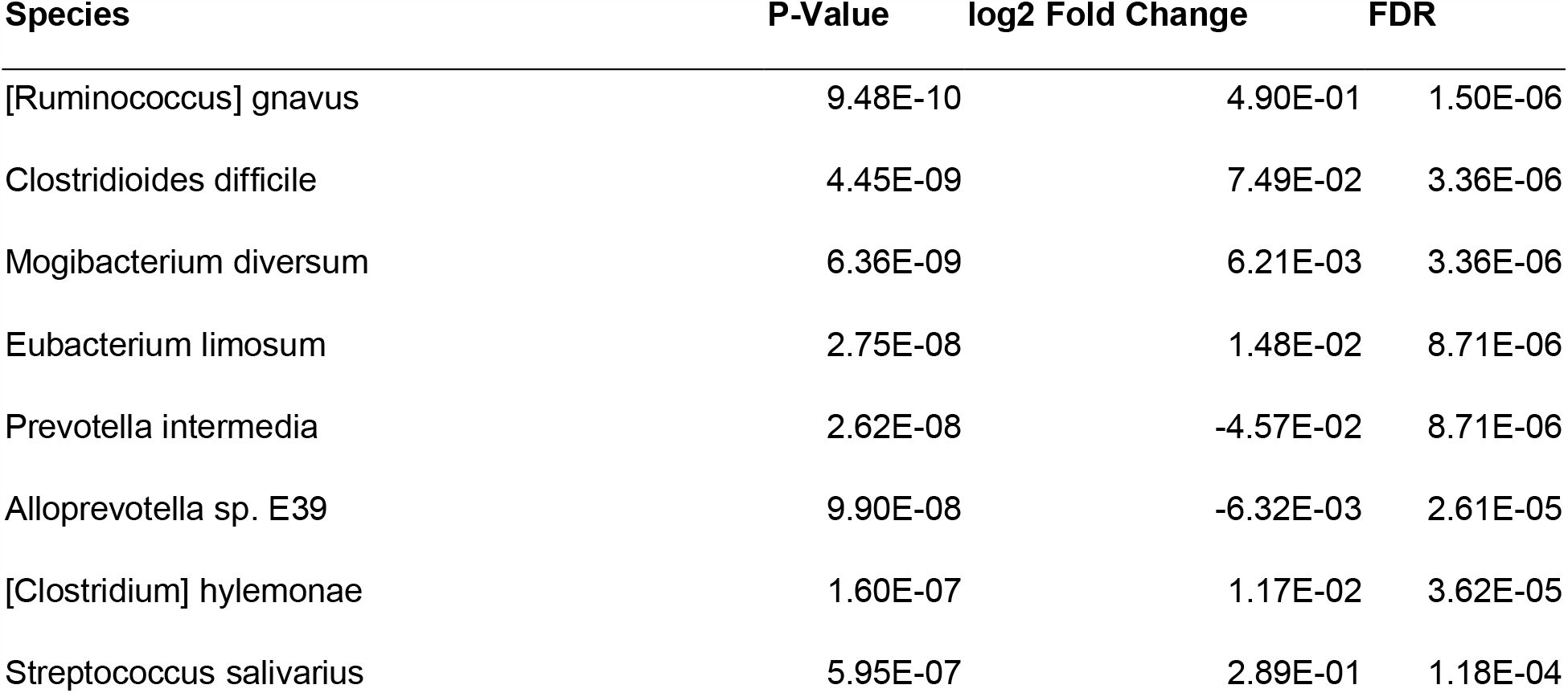
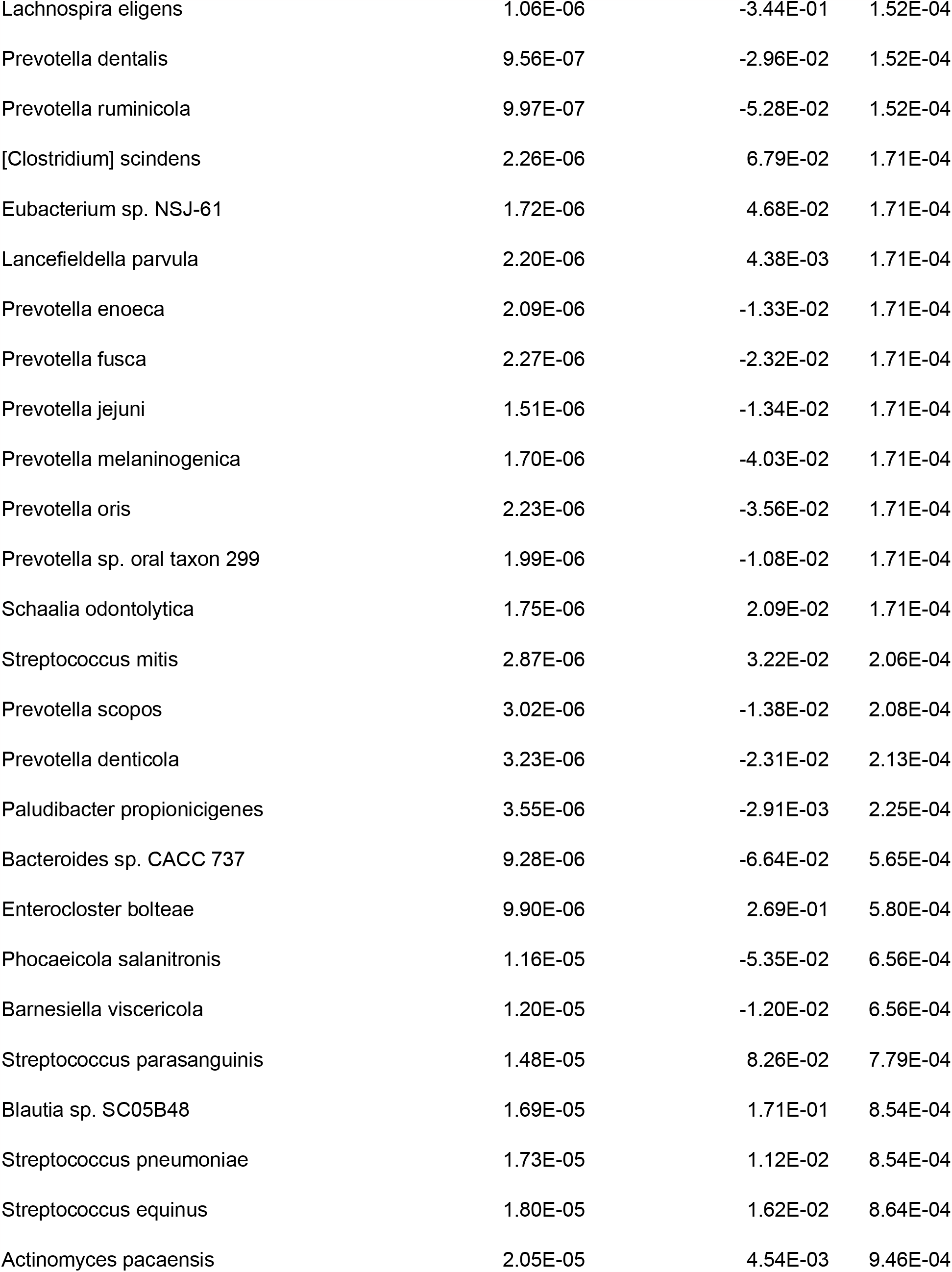
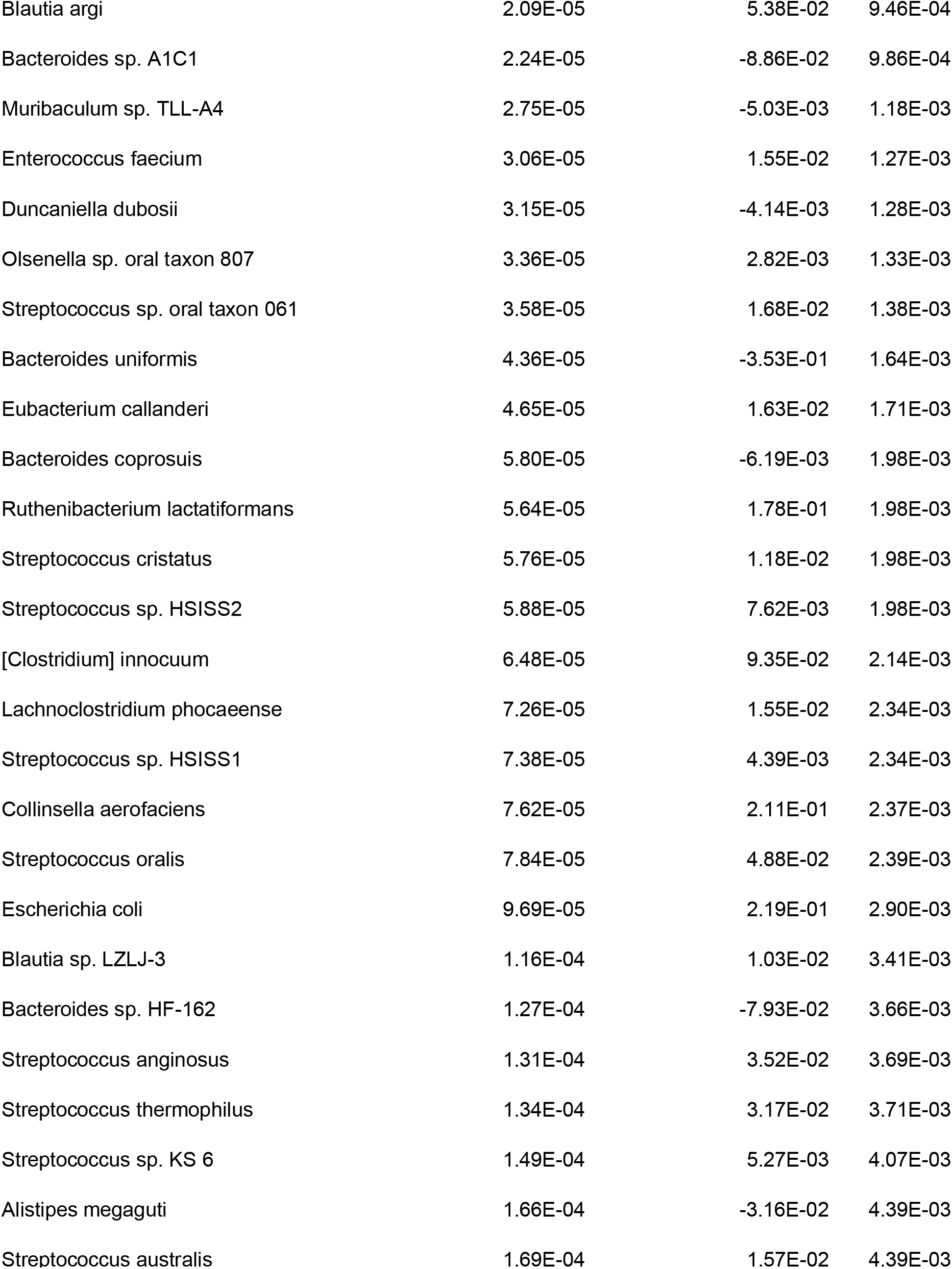
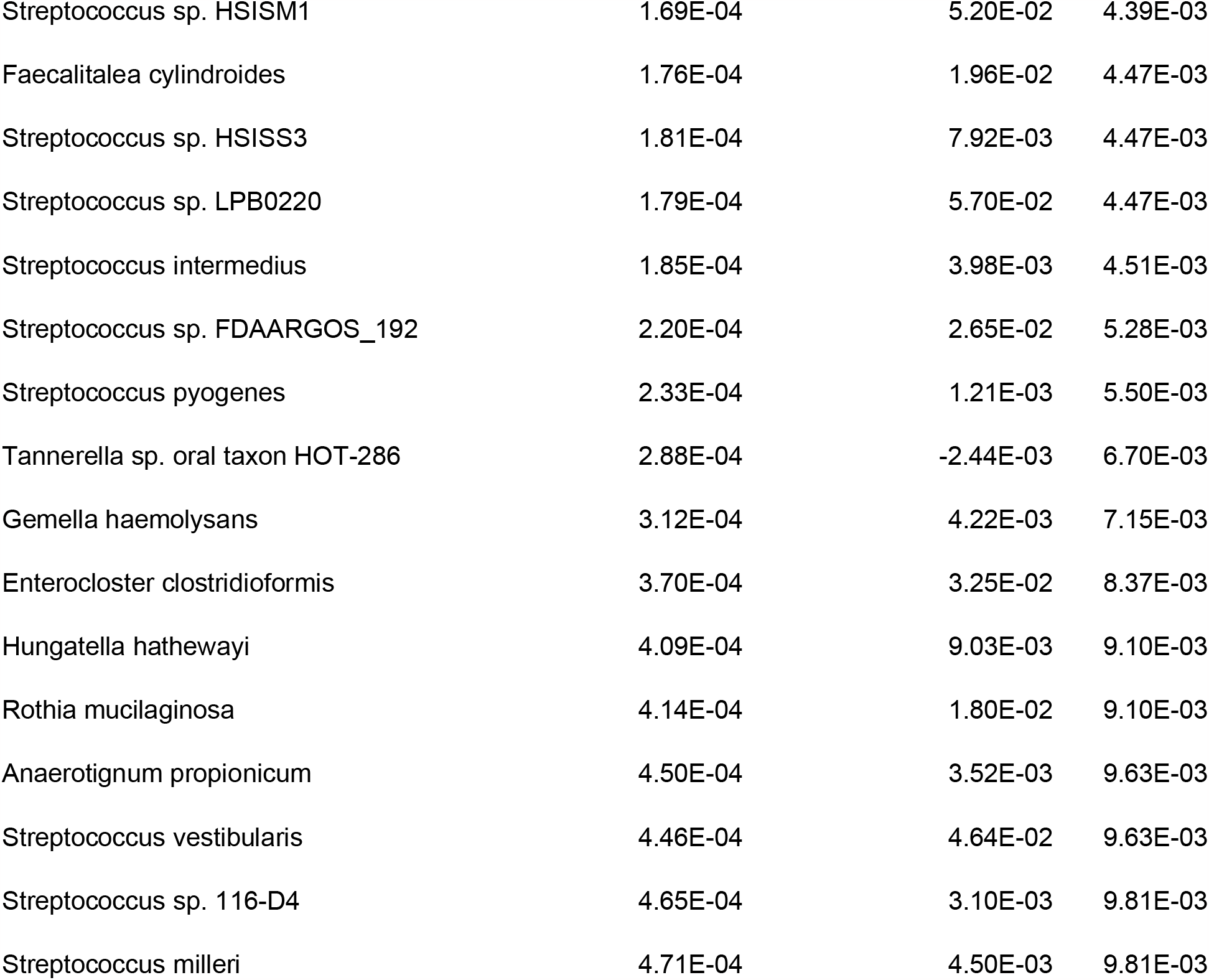
Significant Differentially Abundant Species Identified using a Linear Model filtered at 1% FDR.

**Figure 3:**
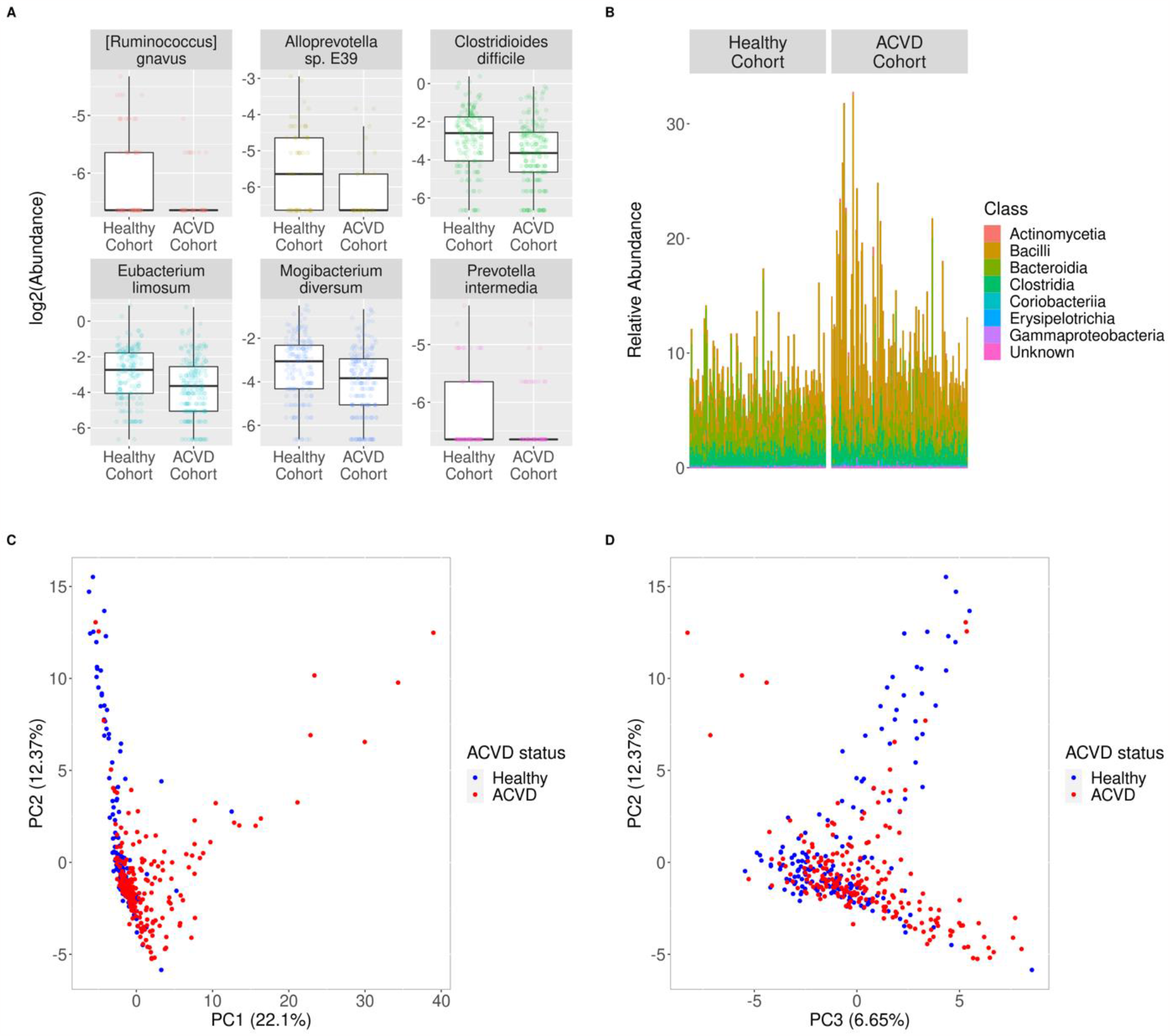
Exploration of Differentially Abundant Microbial Features in Healthy and ACVD Positive Subjects. **(A)** Examples of differentially abundant species between ACVD positive and healthy subjects. Differentially abundant species were found using a linear model with an FDR of 0.01 (n=384 total subjects). **(B)** Taxa abundance profiles of ACVD positive and healthy subjects, at the class level. Abundance profiles are filtered to show just those taxa that are differentially abundant between ACVD positive and healthy subjects (n=384 total subjects). **(C-D)** PCA performed on differentially abundant species (FDR 0.05) between ACVD positive and healthy subjects (n=379, filtered to have Sex label).

We additionally explored the relationship between 3 clinical variables and ACVD status: age, body mass index (BMI), and sex. These clinical parameters were chosen as they had few missing values across samples (n_missing_ = 7, 72, 5 for age, BMI, and sex, respectively across 384 samples) and are easy to measure without additional assays being required. Neither age (p = 0.47, two-sided t-test) nor BMI (p = 0.88, two-sided t-test) showed significant association with ACVD status. In contrast, sex showed significant association with ACVD status (p = 7.75 e-12, fisher’s exact test, Table 2). While this exploration of clinical data does ‘peek’ at test data when looking for associations, we do not expect this to be an issue as it is already known that males suffer a higher risk of atherosclerosis ^16^ and the model itself does not peek at validation or test data during fitting.

### Model Performance

We decided to model ACVD status as a function of position in microbiome PCA space and biological sex (379 samples used due to 5 samples missing sex). Across 10 splits, we obtained a mean AUROC of 0.826 and a mean AUCPR of 0.87 (Figure 4). A full description of the model is provided in the *Online Methods* section and is illustrated in Figure 1.

**Figure 4:**
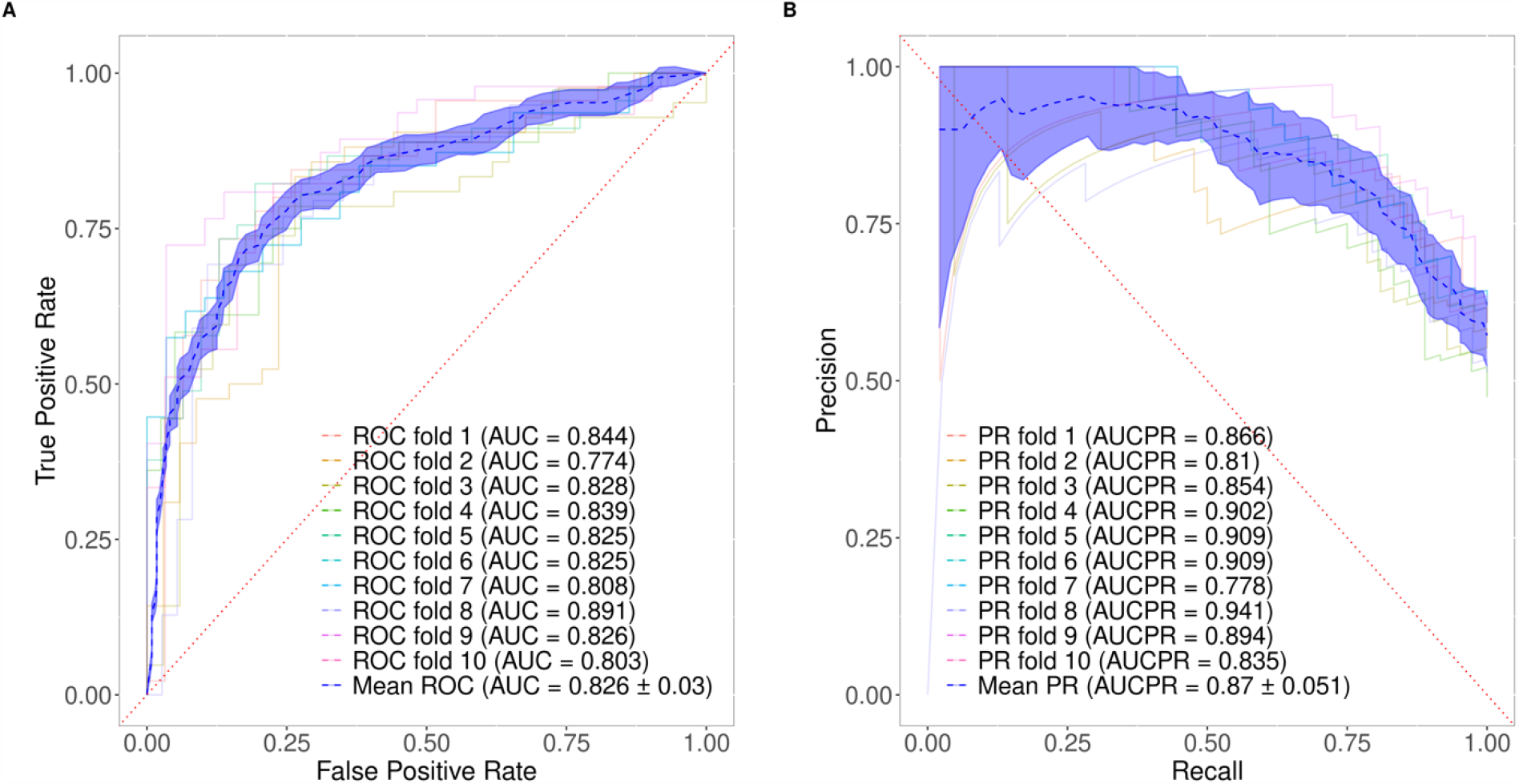
Model Performance with Internal Validation. **(A)** Test AUROC for 10 random splits of data. **(B)** Test AUCPR for 10 splits of data (splits same as in **(A)**).

## Discussion

Atherosclerotic cardiovascular disease presents an enormous cost in human lives and financial resources that is only expected to grow in the coming years. Existing methods for detection are either very accurate in their diagnoses but expensive, or lower cost but with a much lower ability to stratify healthy and diseased individuals. Here, we have shown that shallow depth metagenomic sequencing may provide a viable front-line diagnostic tool towards prioritizing patients for accurate diagnoses at a low cost (Figure 5).

*In-silico* experiments validate the notion of information content preservation as low as 50K reads per sample in terms of species detection and abundance, consistent with current literature.^12^ At a subsampled depth of 50K reads, we observed differentially abundant bacterial taxa in ACVD patients which are consistent findings with previous reports ^10,15,17,18^. In particular, we found a variety of *Streptococcus* species (Table 3) enriched in ACVD patients, species that are associated with increased blood pressure and TMA-lyase.^15^ In addition, we also identified gut microbial species that have been associated with heart failure, *Escherichia coli*, which produce indoxyl sulfate metabolites which induce pro-inflammatory and pro-oxidate effects on cardiomyocytes and cardiac fibroblasts. ^17,19^ Interestingly, there are a few *Bacteroides* that show a low abundance in ACVD patients (Figure S2, Table 4) which has previously been reported and it has been hypothesized that regulation of *Bacteroides spp*. may be important in regulating ACVD progression due to the role of *Bacteroides spp*. in maintaining a healthy gut. ^17^

**Table 4.** Significant Deferentially Abundant Bacterioide Species Identified using a Linear Model filtered at 1% FDR.

| Species | P-Value | log2 Fold Change | FDR |
| --- | --- | --- | --- |
| Bacteroides coprosuis | 5.80E-05 | -6.19E-03 | 1.98E-03 |
| Bacteroides sp. A1C1 | 2.24E-05 | -8.86E-02 | 9.86E-04 |
| Bacteroides sp. CACC 737 | 9.28E-06 | -6.64E-02 | 5.65E-04 |
| Bacteroides sp. HF-162 | 1.27E-04 | -7.93E-02 | 3.66E-03 |
| Bacteroides uniformis | 4.36E-05 | -3.53E-01 | 1.64E-03 |

Given that there is evidently a microbiome ACVD related signature in the shallow depth metagenomics data, we utilize taxa frequency composition information from the sequencing data in conjunction with the patient sex clinical parameter to develop a predictive model of atherosclerotic cardiovascular disease status. We expect that the collection of more patient samples coupled with extensive clinical parameters would improve model performance. Overall, we have demonstrated through *in-silico* subsampling experiments using publicly available data that shallow depth metagenomics sequencing may provide an effective and relatively inexpensive method of diagnosing ACVD status.

## Online Methods

### Dataset

Raw metagenome sequencing data (Illumina HiSeq 2000 paired end) was retrieved from the European Bioinformatics Institute (EBI) public database under accession ERP023788, which was originally acquired by Jie, et. al.^15.^ Of the original 405 Han Chinese subject samples, only 383 samples were retrieved due errors downloading the other 22 samples from the database. The 383 samples includes 170 control subjects (101 females and 68 males) and 214 ACVD subjects (53 females and 157 males). The original study’s inclusion criteria for ACVD diagnosis includes individuals with ≥50% stenosis in single or multiple vessels.^15^ A complete list of samples downloaded and associated clinical metadata used for this study is provided in Table 1.

**Table 1.** Run Accession and Corresponding Available Clinical Parameters. Table 1 is included in a separate document.

### Data Processing

Raw sequencing data was downloaded from the SRA. We then subsampled data using seqtk (v 1.3) ^20^, and cleaned using kneaddata (v 0.7.4) ^21^ using hg37dec_v0.1.4.bt2 as contaminant reference database and trimmomatic (v3.7) for read trimming and quality control, with default options for trimmomatic. Following this, relative abundance estimates were generated using Kraken2 (v 2.1.2) ^22^. For Kraken2, we used the standard Kraken2 bacterial database, downloaded in June 2021.

### Subsampling Experiment

All samples were subsampled at varying depths, ranging from 500 reads to 10 million reads. For all depths below 10 million reads, we determined species abundance profiles, and calculated the total number of features for each sample as well as several overlap metrics to assess overlap of detected features with those detected at 10 million depth.

In order to assess if the subsampling results at 10K depth could be consistently reproduced, we obtained 6 10K depth subsamples on the same set of samples and assessed the number and overlap of detected features across the different seeds.

### Model Definition and Training

The model takes as input biological sex and a relative abundance profile (a_rel_) of bacterial species generated by Kraken2 (see *Data Processing*). During training, differentially abundant features between each class (ACVD positive and negative) are determined with 2 one-sided t-tests, and filtered at an FDR of 0.05. a_rel_ is filtered for the differentially abundant features for each class. Filtered data is then standard normalized and undergoes a PCA transformation (3 PCs). PCA transformed data is then concatenated with biological sex (1 for male, 0 for female), and the concatenated vector is fed to a logistic regression classifier (elastic net model, l1 ratio fixed at 0.5), outputting a label of 1 for ACVD positive and 0 for ACVD negative. When the fit model is queried with new inputs, the feature filtration takes place without new differential abundance determination, and the parameters fit for standard normalization, PCA, and logistic regression are also fixed.

Training is done as follows: 1) split data into train and test set (80:20 split) 2) train data with 5 fold cross validation, refitting with the optimal regularization strength for logistic regression (range for hyperparameter C is exp(-10) to exp(10), with exp(x) increasing in increments of x += 1). 10 random train test splits were performed to assess the distribution of model performance with respect to the sampling distribution.

Software used for the machine learning pipeline include scikit-learn (v1.0.2), scipy (v1.7.3), pandas (v1.3.5), and numpy (v1.21.5)

### Other Notes on Software

Software and versions: Python (v3.7) (anaconda); openjdk (v10.0.2); R (v4.0)

## Contributions

JCS, OW and SD designed the project. JCS and OW implemented data processing and prediction pipeline and performed analyses. JCS, OW and SD wrote the manuscript. The authors have declared no conflict of interest.

## Acknowledgements

We would like to thank Dr. Michael G. Surette and Frank Shannon for insightful discussions and revisions to the manuscript.

## Supplementary Figures

**Supplementary Figure 1:**
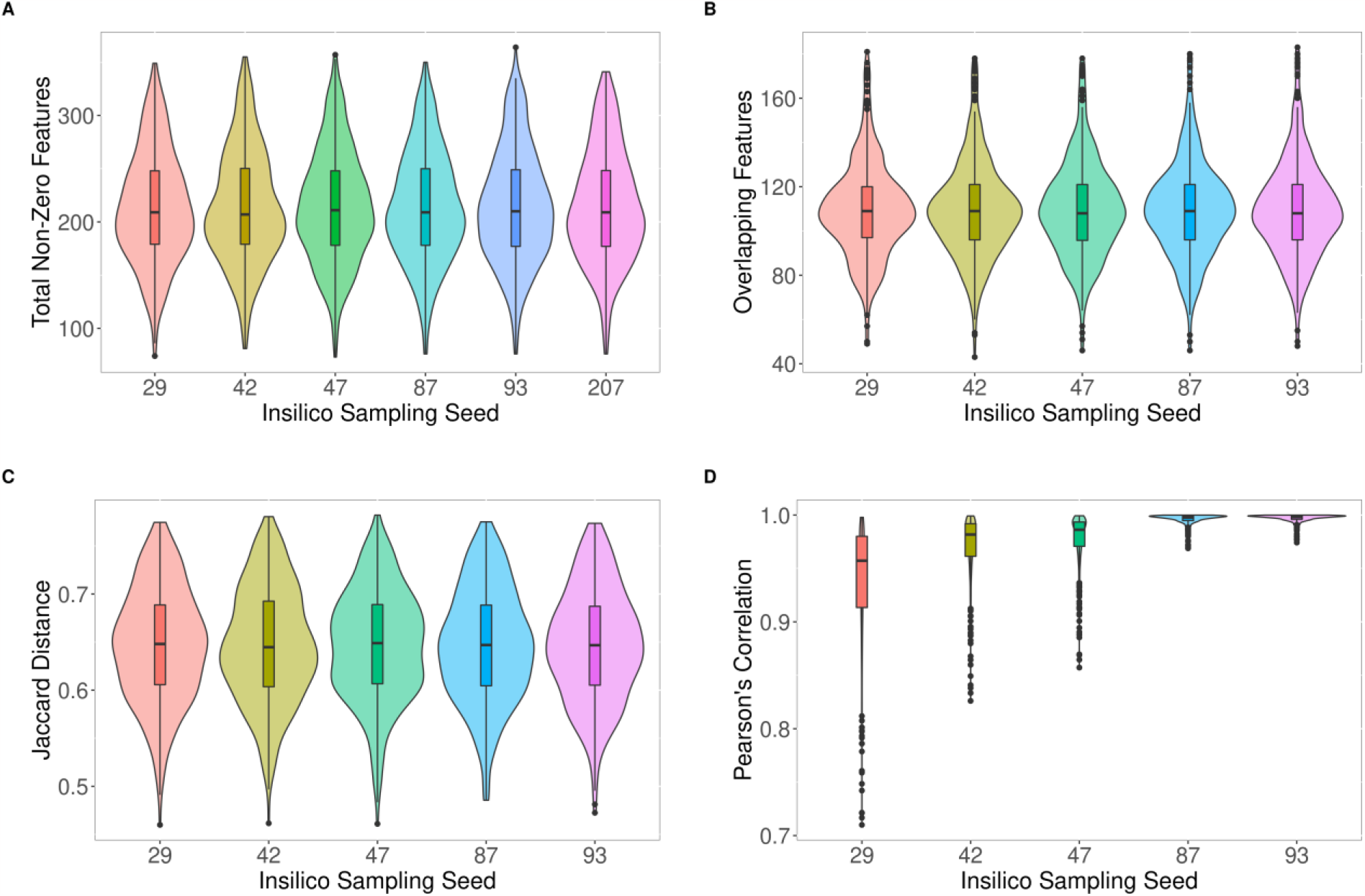
Seed Variability in Feature Detection and Abundance in Microbiome Sequencing Data. **(A)** Total nonzero features detected per sample, by seed. Briefly, we subsample data at 10K depth using a different seed for a random number generator, which will allow us to obtain a different randomly selected reads for each subsample in a reproducible manner **(B-D):** Overlap in feature set, Jaccard distance in feature set, and correlation of overlapping features between identical samples in various seeds when a given seed is compared to seed 207. N = 384 samples.

**Supplementary Figure 2:**
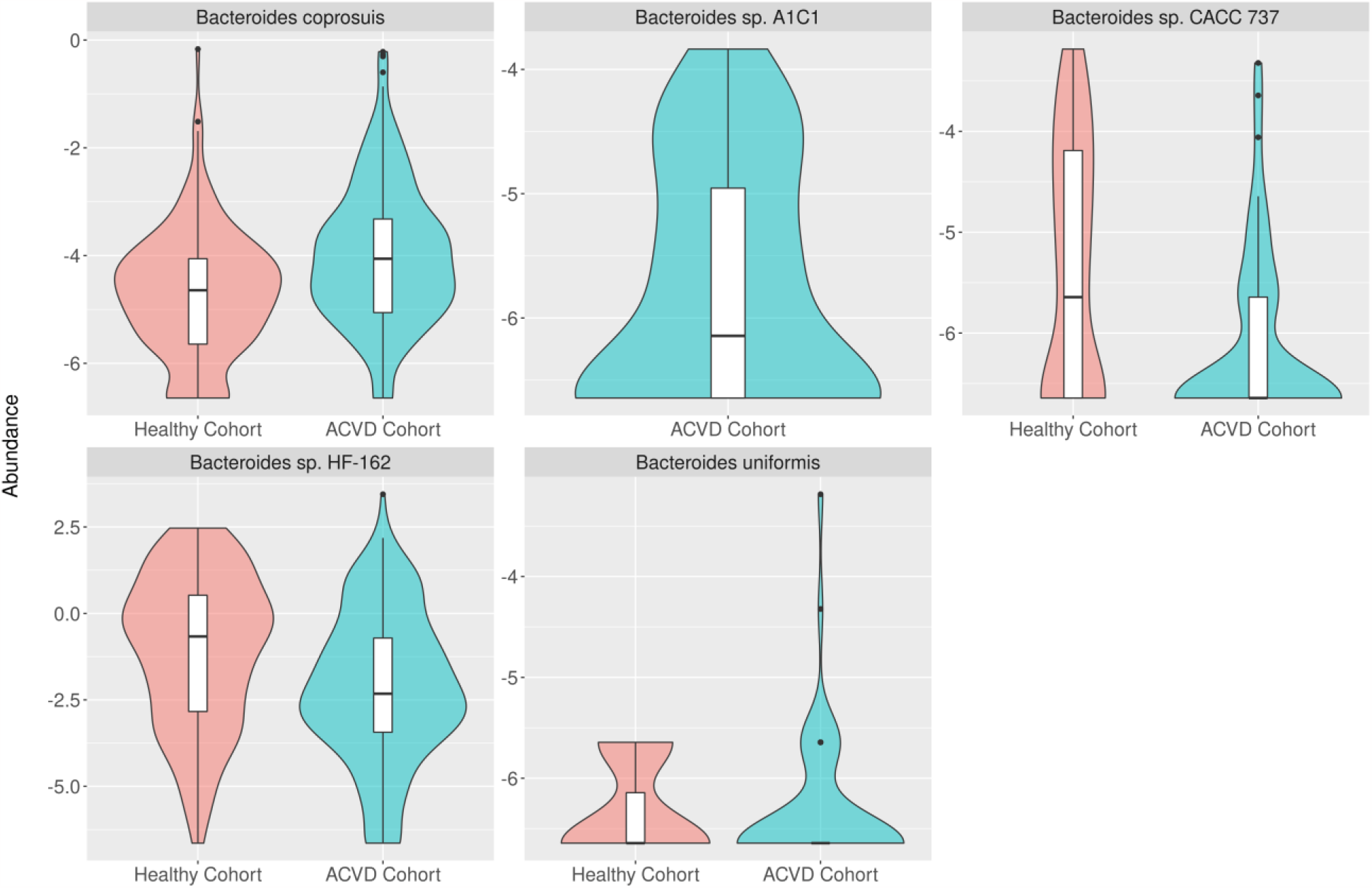
Distribution of Significantly Differentially Expressed Bacteroides species Abundance in ACVD Patients and Healthy Subjects. Distribution of significantly differentially expressed *Bacteroides* species (5 of 21 total species) identified as being differentially abundant at 1% FDR in ACVD vs Healthy patients.

## References

1. Khan T. Cardiovascular diseases. Who.int Web site. https://www.who.int/health-topics/cardiovascular-diseases#tab=tab_1. Updated 2021. Accessed 01 September, 2021.

2. Heart disease facts | cdc.gov. Centers for Disease Control and Prevention Web site. https://www.cdc.gov/heartdisease/facts.htm. Updated 2021. Accessed 01 September, 2021.

3. Roth GA, Mensah GA, Johnson CO, et al. Global burden of cardiovascular diseases and risk factors, 1990–2019: Update from the GBD 2019 study. Journal of the American College of Cardiology. 2020;76(25):2982–3021. https://www.sciencedirect.com/science/article/pii/S0735109720377755. Accessed Sep 1, 2021. doi: 10.1016/j.jacc.2020.11.010.

4. Heidenreich PA, Trogdon JG, Khavjou OA, et al. Forecasting the future of cardiovascular disease in the united states. Circulation. 2011;123(8):933–944. https://www.ahajournals.org/doi/full/10.1161/CIR.0b013e31820a55f5 https://www.ahajournals.org/doi/full/10.1161/CIR.0b013e31820a55f5. doi: 10.1161/CIR.0b013e31820a55f5.

5. Tran Dat. T, Palfrey D, Welsh R. The healthcare cost burden in adults with high risk for cardiovascular disease. PharmacoEconomics - Open. 2021;5(3):425–435. https://link.springer.com/article/10.1007%2Fs41669-021-00257-8. doi: 10.1007/s41669-021-00257-8.

6. Leal J, Luengo-Fernández R, Gray A, Petersen S, Rayner M. Economic burden of cardiovascular diseases in the enlarged european union. European Heart Journal. 2006;27(13):1610–1619. https://doi.org/10.1093/eurheartj/ehi733. Accessed Sep 1, 2021. doi: 10.1093/eurheartj/ehi733.

7. Wielgosz A, Arango M, Bancej C, et al. Tracking heart disease & stroke in canada. Public Health Agency of Canada. 2009:10–13.

8. Lew SQ, Radhakrishnan J. Chronic kidney disease and gastrointestinal disorders. Chronic Renal Disease. 2020:521–539. doi: 10.1016/b978-0-12-815876-0.00033-4.

9. Cresci GAM, Izzo K, Adult Short Bowel Syndrome. Gut microbiome. . 2019:45–54. doi: 10.1016/b978-0-12-814330-8.00004-4.

10. Kazemian N, Mahmoudi M, Halperin F, Wu JC, Pakpour S. Gut microbiota and cardiovascular disease: Opportunities and challenges. Microbiome. 2020;8(36). doi: 10.1186/s40168-020-00821-0.

11. Gao B, Chi L, Zhu Y, et al. An introduction to next generation sequencing bioinformatic analysis in gut microbiome studies. Biomolecules. 2021;11(4). doi: 10.3390/biom11040530.

12. Hillmann B, Al-Ghalith GA, Shields-Cutler RR, et al. Evaluating the information content of shallow shotgun metagenomics. mSystems. 2018;3(6). Accessed Sep 8, 2021. doi: 10.1128/mSystems.00069-18.

13. Davenport T, Kalakota R. The potential for artificial intelligence in healthcare. Future Healthc J. 2019;6(2):94–98. https://www.ncbi.nlm.nih.gov/pmc/articles/PMC6616181/. Accessed Sep 8, 2021. doi: 10.7861/futurehosp.6-2-94.

14. Mathur P, Srivastava S, Xu X, Mehta JL. Artificial intelligence, machine learning, and cardiovascular disease. Clin Med Insights Cardiol. 2020;14. https://www.ncbi.nlm.nih.gov/pmc/articles/PMC7485162/. Accessed Sep 8, 2021. doi:10.1177/1179546820927404.

15. Jie Z, Xia H, Zhong S, et al. The gut microbiome in atherosclerotic cardiovascular disease. Nat Commun. 2017;8(1):1–12. https://www.nature.com/articles/s41467-017-00900-1. Accessed Sep 9, 2021. doi: 10.1038/s41467-017-00900-1.

16. Gao Z, Chen Z, Sun A, Deng X. Gender differences in cardiovascular disease. Medicine in Novel Technology and Devices. 2019;4:100025. https://www.sciencedirect.com/science/article/pii/S2590093519300256. Accessed Oct 3, 2021. doi: 10.1016/j.medntd.2019.100025.

17. Yoshida N, Yamashita T, Hirata K. Gut microbiome and cardiovascular diseases. Diseases. 2018;6(3). https://www.ncbi.nlm.nih.gov/pmc/articles/PMC6164700/. Accessed Sep 20, 2021. doi: 10.3390/diseases6030056.

18. Li J, Li Y, Zhou Y, Wang C, Wu B, Wan J. Actinomyces and alimentary tract diseases: A review of its biological functions and pathology. BioMed Research International. 2018;2018:e3820215. https://www.hindawi.com/journals/bmri/2018/3820215/. Accessed Sep 20, 2021. doi: 10.1155/2018/3820215.

19. Jaglin M, Rhimi M, Philippe C, et al. Indole, a signaling molecule produced by the gut microbiota, negatively impacts emotional behaviors in rats. Front Neurosci. 2018;0. https://www.frontiersin.org/articles/10.3389/fnins.2018.00216/full. Accessed Oct 1, 2021. doi: 10.3389/fnins.2018.00216.

20. Li H. Seqtk, toolkit for processing sequences in FASTA/Q formats. https://github.com/lh3/seqtk.

21. Quality control tool on metagenomic and metatranscriptomic sequencing data, especially data from microbiome experiments. https://github.com/biobakery/kneaddata.

22. Wood DE, Lu J, Langmead B. Improved metagenomic analysis with kraken 2. Genome Biol. 2019;20(1):257. https://doi.org/10.1186/s13059-019-1891-0. doi: 10.1186/s13059-019-1891-0.

